# A standardized and reproducible workflow for membrane glass slides in routine histology and spatial proteomics

**DOI:** 10.1101/2023.02.20.529255

**Authors:** Thierry M. Nordmann, Lisa Schweizer, Andreas Metousis, Marvin Thielert, Edwin Rodriguez, Lise Mette Rahbek-Gjerdrum, Pia-Charlotte Stadler, Michael Bzorek, Andreas Mund, Florian A. Rosenberger, Matthias Mann

**Author notes:** Equal contribution (Thierry M. Nordmann, Lisa Schweizer).

## Abstract

Defining the molecular phenotype of single cells *in-situ* is essential for understanding tissue heterogeneity in health and disease. Powerful imaging technologies have recently been joined by spatial omics technologies, promising unparalleled insights into the molecular landscape of biological samples. One approach involves laser microdissection in combination with membrane glass slides for the isolation of single cells from specific anatomical regions for further analysis by spatial omics. However, so far this is not fully compatible with automated staining platforms and routine histology procedures such as heat-induced epitope retrieval, limiting reproducibility, throughput and integration of advanced staining procedures. This study describes a robust workflow for routine use of glass membrane slides, allowing precise extraction of tissue in combination with automated and multicolor immunofluorescence staining. The key advance is the addition of glycerol to standard heat-induced epitope retrieval protocol, preventing membrane distortion while preserving antigen retrieval properties. Importantly, we show that glycerol is fully compatible with mass-spectrometry based proteomics and does not affect proteome depth or quality. Further, we enable single focal plane imaging by removing remaining trapped air pockets with an incision. We demonstrate our workflow using the recently introduced Deep Visual Proteomics technology on the single-cell type analysis of adjacent suprabasal and basal keratinocytes of human skin. Our protocol extends the utility of membrane glass slides and enables much more robust integration with routine histology procedures, high-throughput multiplexed imaging and sophisticated downstream spatial omics technologies.

## INTRODUCTION

Preservation of spatial context is essential for spatial omics technologies, such as genomics, transcriptomics and proteomics amongst others [1,2]. Laser microdissection (LMD) enables accurate extraction of tissue and single cells from defined anatomical regions, thereby preserving the all-important spatial information [3]. Furthermore, LMD as a single technique is compatible with many downstream omics technologies, providing the opportunity for multiomics analysis from the same sample [4,5]. Glass or metal frame slides coated with a plastic membrane are essential for precise LMD and optimal tissue extraction, in particular when attempting to retrieve anatomical structures at cellular or even subcellular level. There are different types of membranes including polyethylene naphthalate (PEN), polyphenylene sulfide (PPS), polyethylene terephthalate (PET), polyester (POL) or fluocarbon (FLUO), each with unique physical and imaging properties [6]. Despite this variety, the incompatibility of membrane slides with routine histology strategies such as standard heat-induced epitope retrieval (HIER) has been a long-standing problem in spatial omics technologies [7]. Specifically, multiplex immunofluorescence (mIF) can visualize numerous cell types, adding an important layer for omics analyses in a single tissue, but is hampered by the absence of robust multiplexed staining on membrane slides [8,9]. In addition, the majority of studies that require membrane slides employ manual staining techniques, thereby not leveraging the advantage of reproducible and high-throughput staining workflows. This manuscript proposes a solution to these challenges by introducing a protocol for handling of glass membrane slides for routine histological procedures and multiplex IF. Our protocol processes membrane glass slides in high throughput for LMD, without altering or affecting downstream proteomic analysis. Finally, we show that this workflow enables highly precise microdissection at the single cell level while preserving all spatial information and profile these cells using our recently published Deep Visual Proteomics (DVP) [10].

## MATERIAL and METHODS

### Tissue samples

Formalin fixed and paraffin embedded (FFPE) human tissue samples were collected according to standard operating procedures. In more detail, skin specimens were stored in 5% formalin for 24 - 48 hours at room temperature (RT), trimmed and placed in embedding cassettes prior to automated processing (Tissue-TEK VIP, Sakura). Skin tissue sections were collected following informed consent and ethical approval (EK 22-0343). Tissue specimens for the microarray block were fixed in formalin for 24-72 hours at RT, trimmed and manually embedded in paraffin. All experiments were performed in accordance with the Declaration of Helsinki. Regarding the tissue microarrays, according to Danish Data Protection Act (2018), they are exempted from patient consent and permission from legal authorities for fully anonymized material.

### Tissue sectioning

2 μm PEN membrane slides (MicroDissect GmbH, MDG3P40AK) were pretreated with Vectabond (Biozol, VEC-SP-1800) according to manufacturer’s protocol and dried overnight at RT. FFPE tissue blocks were cooled to -17°C at least 1 hour prior to sectioning. Then, 2.5 μm sections were cut using a rotary microtome and placed onto a water bath at 37°C. Sections were transferred to pretreated PEN membrane slides and dried overnight at 37°C.

### Visual evaluation of membrane stability

Polyethylene naphthalate (PEN) membrane slides were treated identical to the initial experimental procedure of the DVP staining protocol described below. After antigen retrieval, slides were washed twice with ddH20 and imaged using the UV-scan mode of a gel documentation system (Axygen, GD-100) and Canon EOS 5D.

### DAKO staining platform

A small tissue microarray block composed of FFPE skin, appendix, cerebellum, adrenal gland, melanoma, and tonsillar tissue was sectioned and mounted on PEN slides as described above. Next, sections were deparaffinized, rehydrated and loaded wet on the fully automated instrument Omnis (Dako, Glostrup, Denmark) based on Dynamic Gap Staining technology and capillary forces. Sections were subjected to antigen retrieval using citrate buffer pH6 (Tri-sodium citrate dihydrate), Target Retrieval Solution (TRS) pH9 (Dako, S2367) and TRS pH6 (Dako, S2369) with or without 10% Glycerol (Sigma Aldrich/Merck, Darmstadt, Germany, G7757) and heated for 60 minutes at 90°C. Slides were subsequently incubated with anti-CK5 (1:200, Leica Biosystems, clone XM26, Newcastle upon Tyne, Great Britain, NCL-L-CK5) for 30 min at 32°C. After washing and blocking of endogenous peroxidase activity, the reactions were detected/visualized using Envision FLEX+ High pH kit (Dako, GV800+ GV821) and Envision DAB+ Substrate Chromogen System (Dako, GV825) according to the manufacturer’s instructions. Finally, slides were rinsed in water, counterstained with Mayer’s hematoxylin and air-dried prior to mounting.

### Proteomics of bulk tissue sections

Tonsil tissue that was stained as described above was collected from corresponding regions on consecutive slides using laser microdissection. Subsequently, samples were lysed in 300 mM Tris/HCl pH 8.0, 12.5% acetonitrile including 5 mM TCEP and 20 mM CAA for 10 min at 90°C, followed by focused ultrasonication (Covaris, AFA®) and repeated heating for 80 min at 90°C. Samples were then further processed as described previously [11].

### Deep Visual Proteomics

#### Immunofluorescence staining

Mounted slides were heated at 56°C for 20 min and deparaffinized (2x 2 min xylene, 2x 1 min 100% EtOH, 95% EtOH, 75% EtOH, 30% EtOH, and ddH20, respectively). Glycerol-supplemented heat induced epitope-retrieval (G-HIER) was performed using preheated 1x TRS pH9 HIER buffer (DAKO, S2367) / 10% glycerol (v/v; Sigma, G7757) in 50 mL conical tube placed in a water bath at 88°C for 20 min. Subsequent to sample blocking using 5% bovine serum albumin in PBS for 30 min at room temperature, primary antibodies targeting pan-CK (rabbit, DAKO Z0622, dilution 1:100) and KRT10 (mouse, Abcam ab76318, dilution 1:800) were incubated for 1.5 hours at 37°C in a wet staining chamber. Following two wash steps in PBS, an incubation with secondary antibodies against mouse-IgG (A647, Invitrogen A32728, 1:400) and rabbit-IgG (A555, Invitrogen A32732; 1:400) was performed for 1 hour at 37°C in a wet staining chamber, followed by 7 min incubation with SYTOX green Nucleic Acid Stain (Invitrogen S7020; 1:700 in ddH20) at room temperature. Slides were then washed twice in ddH2O and allowed to dry briefly, before puncturing the membrane at the proximal end of the slide using a needle (30G) to eliminate the localized membrane elevation followed by subsequent adhesive sealing. Then, tissue sections were mounted with a cover glass using Slowfade Diamond Antifade Mountant (Invitrogen, S36967). After slide scanning, cover glasses were removed from membrane slides upon imaging by an incubation period of 5-10 min in ddH2O and air-dried at RT.

#### Image analysis and laser microdissection

Artificial intelligence (AI)-guided cell recognition, classification and extraction followed by mass spectrometry-based profiling was performed as described recently [10]. In brief, fluorescence images were acquired on a Zeiss Axioscan Z7 at 20X magnification with a 10% tile overlap and analyzed using the Biology Image Analysis Software (BIAS, Cell Signaling). Keratinocytes were identified with a deep neural network on the basis of pan-cytokeratin. After removal of duplicates at tile-overlapping regions, we used a supervised machine learning approach to differentially classify KRT10^pos^ (e.g. suprabasal) from KRT10^neg^ (e.g. basal) keratinocytes. Contour outlines were then exported along with reference points for image registration. Finally, 700 contours of each group were excised in quadruplicates on a LMD7 (Leica Microsystems) and collected into an underlying 384 well plate.

#### Sample processing and mass spectrometry

Sample replicates were processed following our recently published workflow [10]. We used a liquid handling platform (Agilent Technologies, Bravo) to ensure reproducibility and high-throughput processing of samples. In brief, single-cell shapes were collected at the bottom of each well using centrifugation and vacuum evaporation upon the addition of acetonitrile. Subsequently, cells were lysed in 4 μL 60mM TEAB for 60 min at 95°C, followed by further 60 min at 75°C in 12% (v/v) acetonitrile. Protein digest was performed overnight using 4 ng LysC and 6 ng trypsin in a total sample volume of 7.5 μL, respectively. The enzymatic reaction was quenched in a final concentration of 1% (v/v) trifluoracetic acid (TFA). Subsequently, samples were loaded on Evotips Pure tips following the manufacturer’s instructions.

### LC-MS/MS analysis of bulk and ultra-high sensitivity data

Bulk samples were reconstituted in buffer A* (2% Acetonitrile (ACN)/0.1% formic acid (FA) in LC-MS grade water). MS data was acquired by an EASY nanoLC 1200 (Thermo Fisher Scientific) coupled to a timsTOF Pro2 mass spectrometer (Bruker Daltonics) with a nano-electrospray ion source (CaptiveSpray, Bruker Daltonics). 200 ng of peptides were loaded on a 50 cm in-house packed HPLC column (75 μm inner diameter packed with 1.9 μm ReproSil-Pur C18-AQ silica beads, Dr. Maisch GmbH, Germany). The column temperature was kept at 60 °C by an in-house manufactured oven. Sample analytes were separated using a linear 120 min gradient from 3 to 30% buffer B in 95 min, followed by an increase to 60% for 5 min and to 95% buffer B for 5min, as well as a 5 min wash at 95% buffer B and re-equilibration for 5 min at 5% buffer B (buffer A: 0.1% FA/ddH_2_O; buffer B: 0.1% FA, 80% ACN, and 19.9% ddH_2_O). The flow rate was kept constant at 300 nl/min. The mass spectrometer was operated in dda-PASEF mode as previously described [12]. Briefly, one MS1 scan was followed by ten PASEF MS/MS scans per acquisition cycle. The ion accumulation and ramp time in the dual TIMS analyzer was 100 ms and ion mobility range was set from 1/K0 = 1.6 Vs cm^−2^ to 0.6 Vs cm^−2^. Single charged precursor ions were excluded with a polygon filter (timsControl, version 3.0.20.0, Bruker Daltonics) and precursors for MS/MS were picked at an intensity threshold of 2,500 arbitrary units (a.u.) and re-sequenced until reaching a target value of 20,000 a.u. considering a dynamic exclusion of 40 s elution.

For the high-sensitivity, deep-visual proteomic samples, samples were loaded onto Evotips Pure and measured with the Evosep One LC system (Evosep) coupled to a timsTOF SCP mass spectrometer (Bruker) employing a nano-electrospray ion source (Bruker Daltonics). The Whisper 20 SPD (samples per day) method was used with the Aurora Elite CSI third generation column with 15 cm and 75 μm ID (AUR3-15075C18-CSI, IonOpticks) at 50 °C. The mobile phases comprised 0.1% FA in LC-MS grade water as buffer A and 99.9% ACN/0.1% FA as buffer B. The timsTOF was operated in dia-PASEF mode with variable window widths. Optimal dia-PASEF methods cover the precursor cloud highly efficient in the m/z – ion mobility (IM) plane while providing deep proteome coverage. For method generation with py_diAID, the precursor density distribution in m/z and IM was estimated based on a tryptic 48 high-pH fraction library [13]. We calculated the optimal cycle time based on the chromatographic peak width of 5 ng HeLa single runs. The optimal dia-PASEF method consisted of one MS1 scan followed by twelve dia-PASEF scans with two IM ramps per dia-PASEF scan, covering a m/z range from 300 to 1200 and IM of 0.7 to 1.3 Vs cm^-2^. All other settings were as described above.

### Data processing

Data acquired in dda-PASEF mode were analyzed with MaxQuant (2.0.1.0) using standard settings in reference to UniProt human databases (UP000005640_9606.fasta and UP000005640_9606_additional.fasta) [14]. Oxidation (M) and Acetyl (Protein-N-term) was set as variable and Carbamidomethyl (C) as fixed modification. The maximum number of modifications per peptide was set to 3. Trypsin/P was selected as protease and the maximum number of missed cleavages was set to 2. MS data acquired in dia-PASEF mode were processed in the DIA-NN software (version 1.8.1, [15]) using the same fasta files and standard settings. Search parameters deviated from the standard settings as follows: Peptide length range 7-55, Precursor m/z range 100-1700, Fragment ion m/z range 100-1700, Quantification strategy ‘any LC (high accuracy)’, cross-run normalization ‘global’, and MBR (‘match between runs’) enabled. The report.pg_matrix output file of DIA-NN was used for further data analysis.

### Bioinformatics data analysis

Data analysis was performed in R v4.2.2. Statistical analysis of bulk data was performed with limma v3.52.4. The number of significant hits (FDR < 5%) was corrected for multiple testing within the shown comparison using a Benjamini-Hochberg correction. The theoretical isoelectric point of a protein was calculated with the computePI function of seqinr v4.2-23 for every protein within one protein group. Hydrophobicity was estimated with the hydrophobicity function (scale = “KyteDoolittle”) in the package peptides v2.4.4. Amino acid sequences for Uniprot identifiers were retrieved with the UniProt.ws package v2.36.5. For protein groups (sequences that cannot be distinguished by the underlying peptide identifications), the mean value of individual proteins is presented.

For the analysis of DVP data, differential protein expression was determined as described above. Biological pathway enrichments were performed using the WebGestaltR package v.0.4.4 for an overrepresentation analysis in reference to the ‘Reactome’ database and an FDR threshold of 5% in the background of all identified proteins in the dataset (organism: homo sapiens). Spatial data from xml files was plotted with the package sf v1.0-9 as used previously [16]. For the principal component analysis, data were filtered for 80% valid values in each group and imputed from left-shifted normal distribution (shift = 1.8, scale = 0.3). The FactoMineR v2.6 package was used to perform the PCA analyses. Visual evaluation of membrane stability was performed in Python 3.9.12 using the NumPy, Matplotlib and Scikit-image packages. Intensity values of the acquired images were normalized to the median intensity of each image and then plotted as heatmaps using the ‘viridis’ colormap, indicating intensity values.

## RESULTS and DISCUSSION

In our routine work with PEN membranes, we regularly observed the formation of air pockets at random locations underneath the membrane during the standard heat-induced epitope retrieval (HIER) protocols, indicating incompatibility with staining, scanning, and laser microdissection procedures (Figure 1A, Supplementary Figure 1A). To address this issue, we aimed at developing a robust HIER protocol with respect to membrane compatibility. Building on the observation that glycerol in HIER buffers enhances antigen retrieval, we noticed that the addition of glycerol had a stabilizing effect on membrane integrity [17]. After HIER it eliminated the formation of random membrane irregularities (Figure 1B, Supplementary Figure 1B).

**Figure 1.**
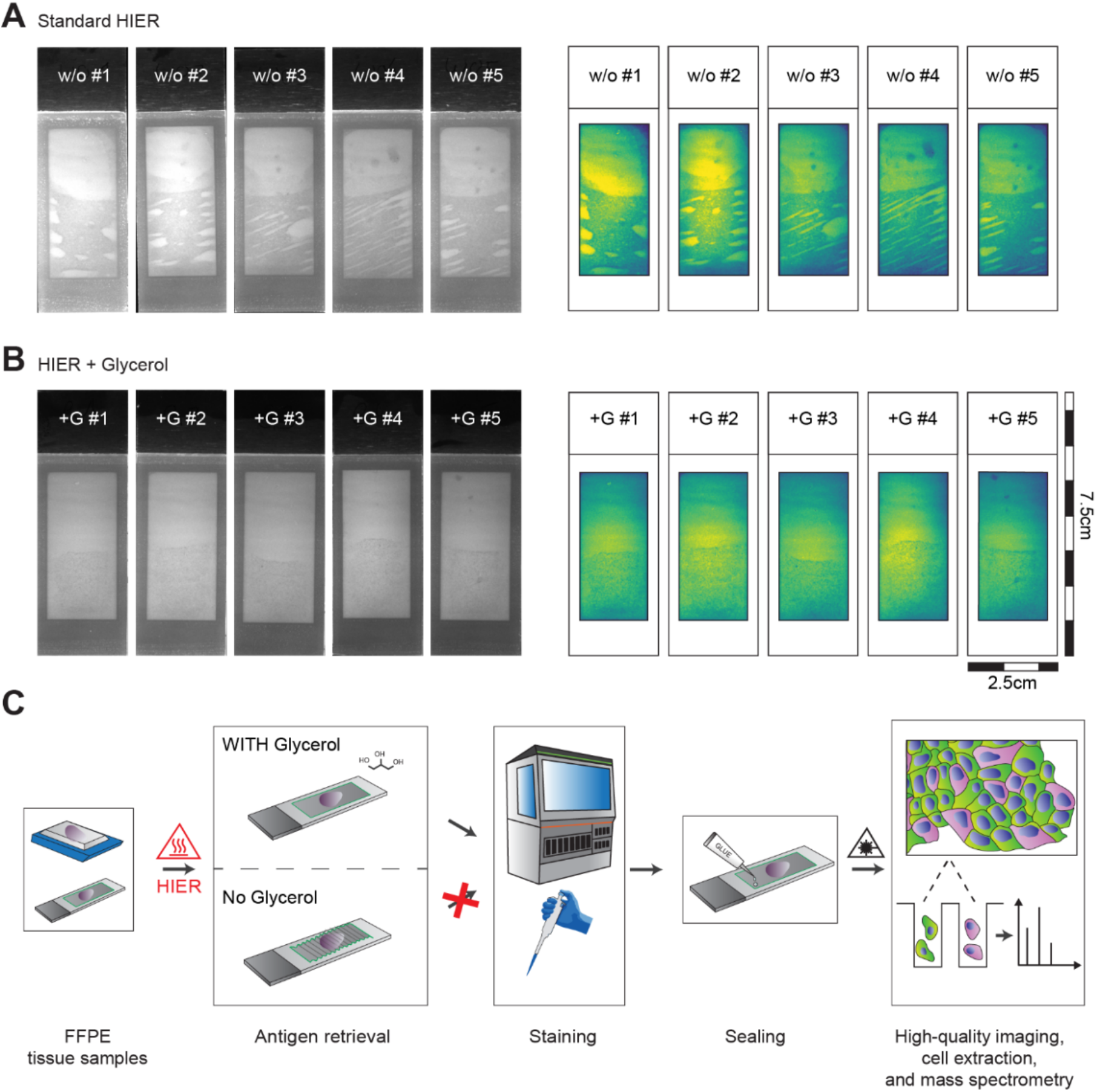
A robust workflow for HIER antigen-retrieval on membrane microscopy slides. (A) Membrane distortion resulting from a standard HIER protocol shown as UV-scans (left) and heatmap of normalized median UV signal intensities across slides. (B) Intact membrane stability using glycerol-supplemented antigen-retrieval in the same arrangement. (C) Workflow schematic for the preparation of LMD-compatible membrane slides. FFPE archival tissue was mounted on 2 μm PEN membrane slides as in routine pathology followed by standard heat-induced epitope-retrieval (HIER) without and with 10% glycerol. While the slides without glycerol were unusable for further staining procedures upon heating in solution, the presence of glycerol stabilized the membrane. Minor accumulation of gas below the membrane were removed using minimal invasive methods followed by glue-based sealing. w/o: without glycerol; +G: HIER buffer supplemented with 10% glycerol.

Following glycerol-supplemented HIER (which we term ‘G-HIER’), tissue sections could readily be processed for staining. However, after staining, multiple imaging planes (z-stacks at 20x resolution) were still needed to capture all tiles of the slide. We found that puncturing the membrane with a 30G syringe and resealing it at its proximal end after completing the staining process, eliminated any remaining surface irregularities and further facilitates high-resolution and rapid whole-slide imaging (Supplementary Figure 1C). As a result, G-HIER allows acquisition in a single focal plane, minimizing the amount of data storage necessary and reducing stitching errors that are detrimental for subsequent laser microdissection.

Having optimized HIER on membrane slides, we tested its compatibility with an automated staining device (Dako Omnis, Agilent) and downstream MS-based proteomics (Fig. 1C). We compared proteome depth and information content of FFPE tonsil tissue samples after antigen retrieval with six different buffer compositions: 10mM citrate buffer at pH 6, and commercially available DAKO buffer at pH 6 and pH 9, with or without addition of 10% glycerol, respectively. To this end, we mounted consecutive, 2.5 μm thick FFPE tissue sections on PEN-membrane slides, performed G-HIER and automated staining against cytokeratin 5 (CK5) using the Dako Omnis platform, followed by laser microdissection and mass spectrometry. We found that, after G-HIER, membrane glass slides were fully compatible with an automated staining device, using the Dako Omnis platform. Proteomic analysis revealed that the number of proteins was almost identical across all buffer conditions, at 5,468 ± 12 proteins (mean ± SD, Figure 2A). In our experience, such proteome depth provides substantial coverage of most biological pathways. Likewise, distribution and median of MS intensity per protein and the coefficient of variation as a measure of variability between replicates were comparable, demonstrating that glycerol has no negative impact (Figure 2B – 2D). Notably, in this comparison, we only found a difference due to the pH of the HIER buffer, while glycerol was not a main driver of sample separation along principal component one and two (Figure 2E,F). We speculated that the pH could affect solubility of proteins depending on their physiochemical properties. Indeed, we found that the theoretical isoelectric point and hydrophobicity, rather than protein length or mass, affected which proteins were retained upon HIER at pH 6 compared to pH 9 (Figure 2G). Although this does not affect G-HIER, it can be taken into consideration when establishing stainings of particular challenging protein targets. Thus, our optimized workflow efficiently combines HIER on membrane slides with downstream proteomics and automated staining techniques, without compromising reproducibility and MS data depth.

**Figure 2.**
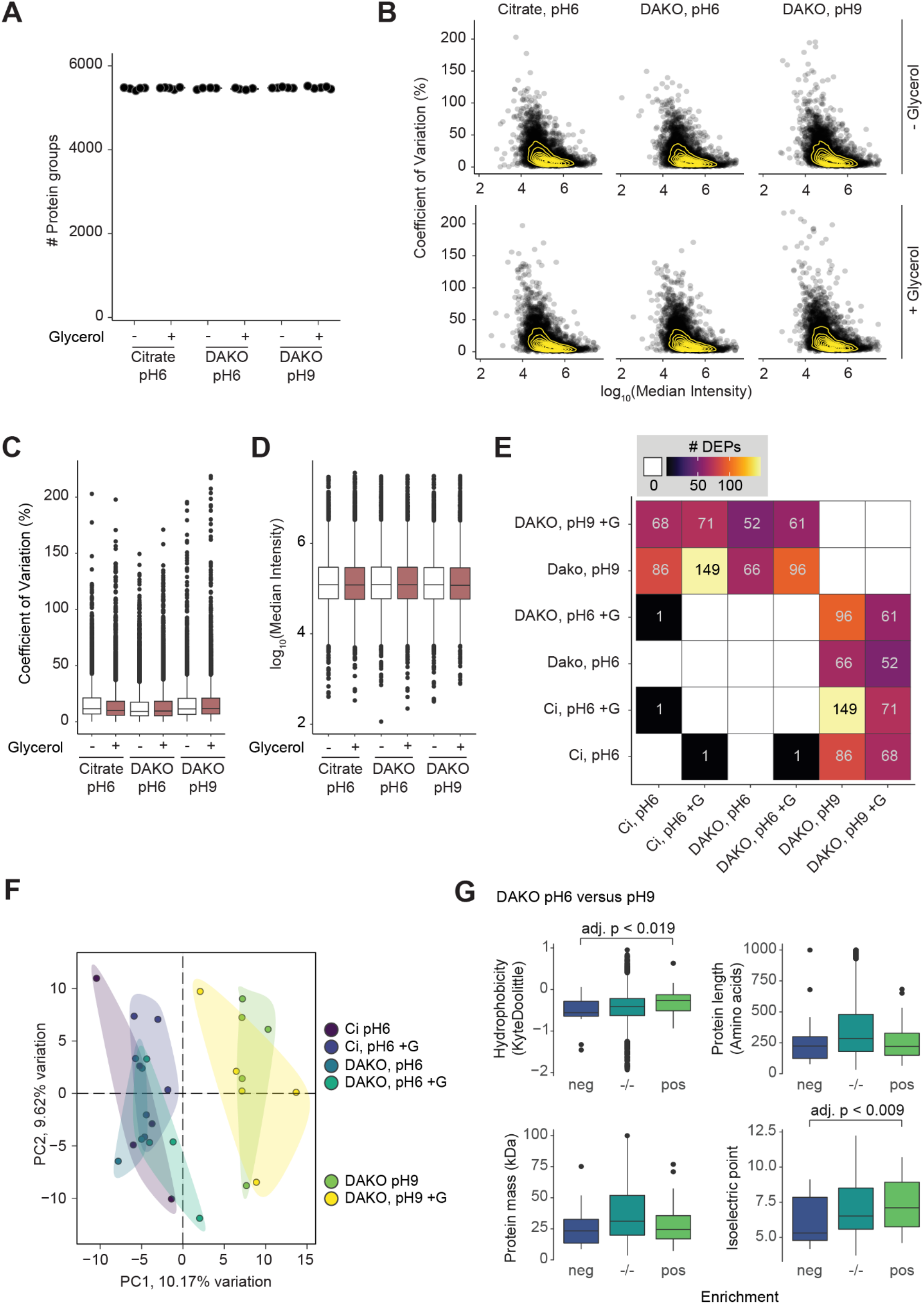
Effect of glycerol in various antigen retrieval solutions on the bulk proteome of tissue slices. (A) Number of proteins detected in laser-dissected tonsil tissue after LC-MS/MS. Points are individual samples, the line represents the median within one group (n > 4). (B) Log-transformed MS signal versus coefficient of variation (%) per solution type. Points are median values of individual proteins per group, density is estimated by yellow lines. (C) Coefficient of variation and (D) log_10_-transformed MS signal as in (B) per group (n > 4). (E) Number of differentially expressed proteins (multiple-testing adjusted p value < 0.05) between indicated comparisons, additionally illustrated with a color gradient. White and empty indicates that no significant hits were found. (F) Principal component analysis of all included samples (n = 28). Colors denote groups with different retrieval solutions. The amount of variation explained per principal component is given with the axis legend. (G) Comparison of proteome composition after heat-induced antigen retrieval with two solutions at pH6 or pH9 (without glycerol). Hydrophobicity and isoelectric point are calculated values. For protein groups, the median value of each contributing protein is shown (n > 4). Boxplots represent median and the 25- and 75-percentile, and whiskers span the 1.5-fold interquartile range. Outliers are shown, if not indicated differently. Ci: Citrate; +G: respective buffer supplemented with 10% glycerol; adj. p.: adjusted p-value; DEP: Differentially expressed proteins.

With a robust protocol in hand, we next aimed to demonstrate its biological relevance by profiling adjacent epidermal cell types of healthy human skin. For this, we integrated G-HIER into the recently developed Deep Visual Proteomics workflow to stain an FFPE tissue specimen by immunofluorescence for pan-cytokeratin and cytokeratin 10 [11]. An AI-based algorithm segmented cells employing the signal of pan-cytokeratin and classified them into basal and suprabasal cells by the presence or absence of cytokeratin 10 (Figure 3A). Laser-assisted cell extraction and subsequent mass spectrometry enabled clear differentiation between the cell types based on the identification and quantification of more than 3,500 proteins across all samples (Figure 3B,C). Between the two cell types, 17% of the proteome was significantly differentially expressed (min. absolute fold change: 1.5, adjusted p-value < 0.05, Figure 3D). Cytokeratin 10 (KRT10), the initial staining target of our study, and KRT1 were upregulated in suprabasal cells, providing positive control. Conversely, common markers of basal keratinocytes such as cytokeratin 15 (KRT15) and basal membrane anchorage fibrils COL4A1 and COL7A1 were highly enriched in the basal cell layer. Summarizing the proteomics results by a biological pathway enrichment analysis, processes such as ‘Keratinization’ and ‘Metabolism of lipids’ reflected the formation of the epidermal barrier in the suprabasal layer, whereas ‘Type I hemidesmosome assembly’ and other cell-cell interactions indicated structural anchoring of cells in the basal layer (Figure 3E). We next took advantage of the integration of spatial and proteomic information to reconstruct the cellular architecture of the skin specimen, mapping the mean protein intensities of KRT1 and KRT15 into the cellular environment (Figure 3F,G). These results demonstrate the robust nature of G-HIER and DVP and how this combination enables the cell-type resolved characterization of human skin by MS-based proteomics while preserving the spatial information.

**Figure 3.**
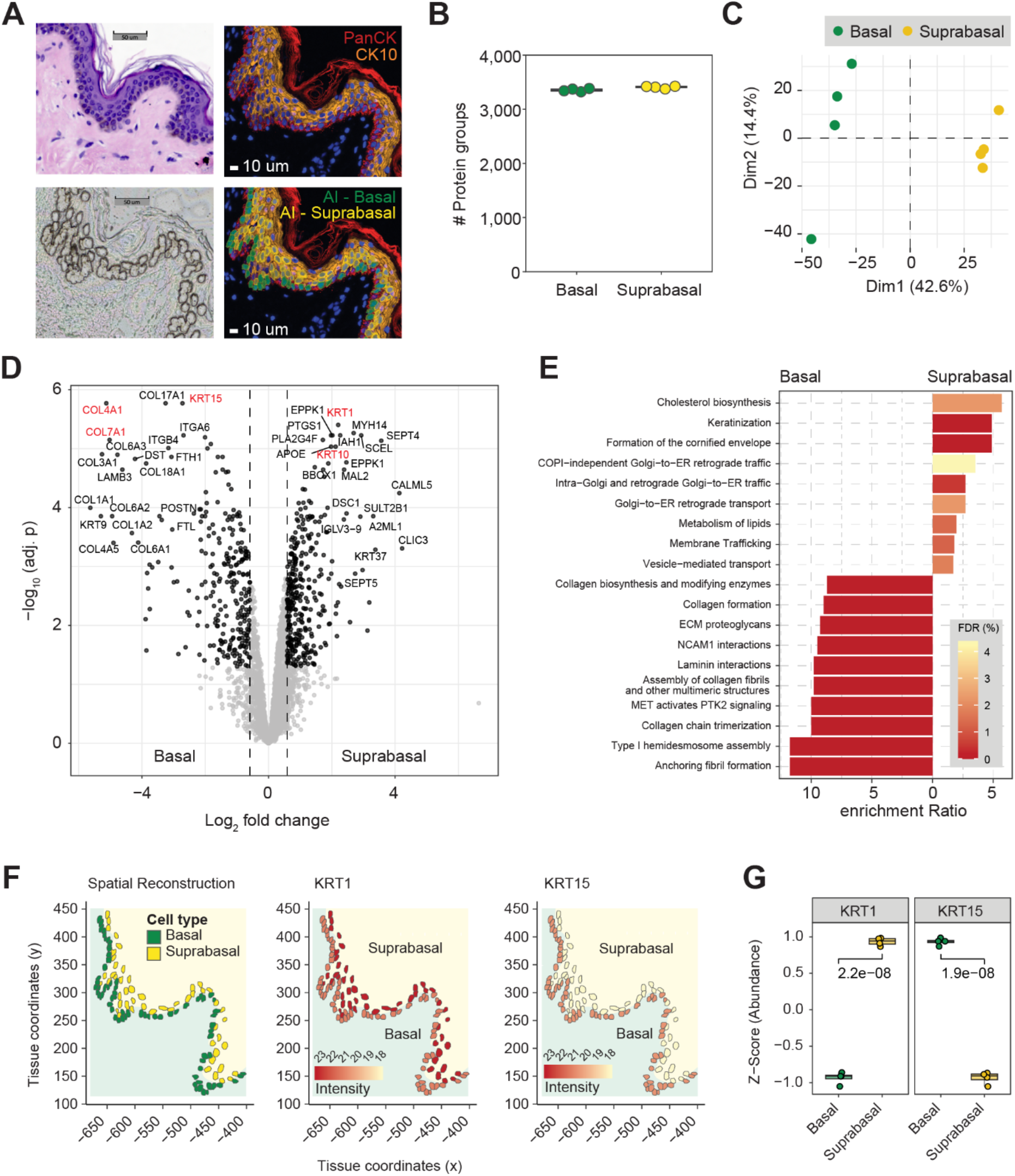
Cell type-resolved spatial proteomics of the human skin. (A) Exemplary region of skin specimen shown in H&E (upper left), IF for PanCK and CK10 (upper right), cell segmentation and classification by artificial intelligence (lower right) and subsequent to cell extraction by laser capture microdissection (lower left). (B) Number of identified protein groups for basal and suprabasal cells of the skin. (C) Principal Component Analysis separating skin cell types in the first two dimensions. (D) Differential protein expression of cell types. Significant proteins (minimum fold change: 1.5, adjusted p-value < 0.05) are highlighted in black, proof-of-principle proteins are shown in red. (E) Biological pathway enrichment analysis based on the ‘pathway Reactome’ database. (F) *In silico* spatial reconstruction of the tissue architecture showing the localization of basal and suprabasal cells in an exemplary tissue region (left panel). Enabled by DVP, mean protein intensities for the representative markers KRT1 (middle panel) and KRT15 (right panel) were mapped onto the tissue architecture. (G) Normalized abundances for the previously shown markers in each cell type. An unpaired student’s t-test was used to annotate the significance for each comparison. adj. p.: adjusted p-value.

In summary, the addition of glycerol to the HIER buffer effectively stabilizes the membrane and enables subsequent high-throughput staining and laser microdissection, which is a crucial step towards robust single-cell or cell type proteomics workflows with extremely sparse material [16]. Our optimized workflow drastically enhances the efficiency of high-throughput LMD workflows and allows more efficient use of precious clinical material, particularly when only a few sections are available for research purposes. Moreover, the addition of glycerol facilitates high-resolution imaging, while preserving antigen retrieval properties, making the use of glass membrane slides more user friendly and as simple as standard glass slides in pathology. In addition, fully automated staining platforms such as the Dako Omnis can be integrated seamlessly to further increase throughput and reproducibility. Of note, evaluation of long-term effects of glycerol on a Dako Omnis instrumentation itself have not been tested. Our workflow renders IF in routine histology fully compatible with the demands of a laser-microdissection workflow on glass membrane slides, which in turn can readily be coupled to DVP or other spatial omics technologies.

## ACKNOWLEDGMENTS

We thank members of the department of Proteomics and Signal Transduction for help and fruitful discussions and especially Katharina Zettl for superbly performing the immunofluorescence staining and Dirk Wischnewski for meticulous sample preparations. Susanne Vondenbusch-Teetz is acknowledged for the scientific implementation of her photography skills. We thank Peter Horvath and Single Cell Technologies Ltd. for their technical support and Fabian Coscia for very valuable feedback.

## AUTHORSHIP CONTRIBUTIONS

The study was conceptualized by F.A.R., T.M.N. and L.S. Experiments were performed by T.M.N., L.S., F.A.R., A.M., M.T., E.R. and M.B. Clinical tissue was collected and prepared by L.M.R.-G., M.B. and P.-C.S. Data were analyzed by L.S., F.A.R., T.M.N., and E.R. The project was supervised by M.M. and A.M. The original manuscript draft was written by T.M.N., F.A.R., L.S. and M.M. All authors read, revised, and approved the manuscript.

## FUNDING

T.M.N. is supported by a Swiss National Science Foundation (SNSF) Early Postdoc Mobility fellowship (P 2ZHP3-199648) and F.A.R. is an EMBO postdoctoral fellow (ALTF 399-2021). This study has further been supported by the Max-Planck Society for Advancement of Science, the Federal Ministry of Education and Research (BMBF) through project CLINSPECT-M, the Chan Zuckerberg Initiative (CZF2019-002448), and by grants from the Novo Nordisk Foundation, Denmark (grant agreements NNF14CC0001 and NNF15CC0001).

## CONFLICT OF INTEREST

M.M. is an indirect investor in Evosep Biosciences. All other authors have no conflicts of interest to declare.

## SUPPLEMENTARY FIGURE

**Supplementary Figure 1.**
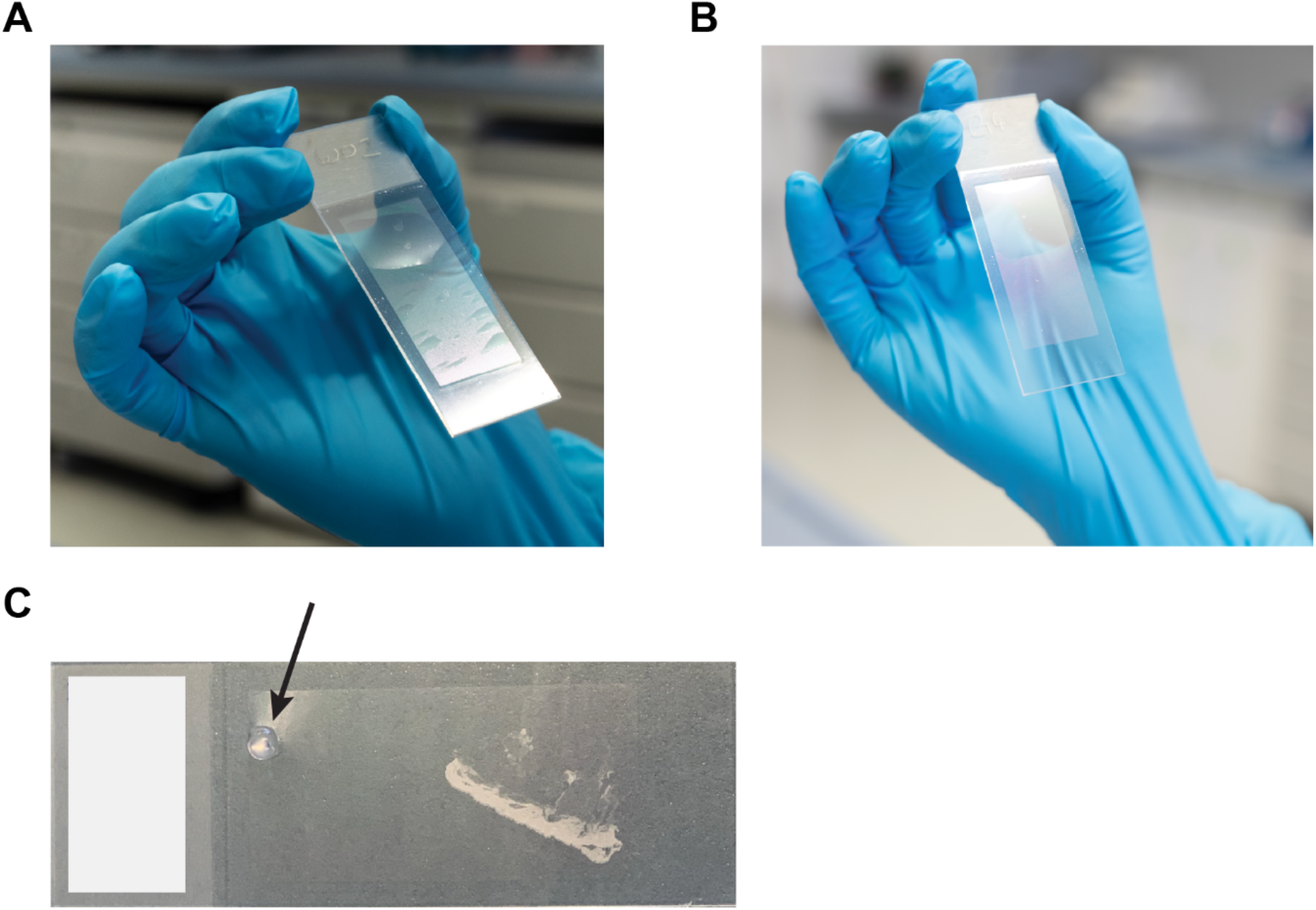
Visible effect of glycerol-assisted HIER on membrane slides. (A) Photography showing the membrane with standard glycerol compared to (B) the addition of 10% glycerol to HIER. (C) Air pockets are removed and sealed enabling single focal-plane imaging.

## Notes

### Competing Interest Statement

M.M. is an indirect investor in Evosep Biosciences. All other authors have no competing interests to declare.

